# Transcutaneous Vagus Nerve Stimulation boosts evidence accumulation during perceptual decision-making

**DOI:** 10.1101/2025.06.27.662007

**Authors:** Shiyong Su, Thomas Vanvoorden, Pierre Le Denmat, Alexandre Zénon, Julie Duque

**Affiliations:** Cognition and Actions Lab, Institute of Neuroscience, UCLouvain, Brussels, Belgium; Department of Brain and Cognition, KU Leuven, Leuven, Belgium; Institut de Neurosciences Cognitives et Intégratives d’Aquitaine, Bordeaux, France

**Keywords:** locus coeruleus, norepinephrine, transcutaneous vagus nerve stimulation, decision-making, random dot motion, drift diffusion model, gain modulation, attention, evidence accumulatio, cognitive control

## Abstract

The locus coeruleus-norepinephrine (LC-NE) system has been implicated in perceptual decision-making, but its causal contribution and underlying mechanisms in humans remain unclear. Here, we used transcutaneous vagus nerve stimulation (tVNS) to modulate LC-NE activity during a random dot motion task, with stimulation delivered at three distinct time points across groups, each targeting different stages of LC-NE engagement during the task. tVNS reliably increased pupil-linked LC-NE activity across all groups. Notably, early stimulation, at a time when LC-NE activity was still at baseline, elicited a more sustained pupil dilation that extended into the decision phase, resulting in comparable pupil responses during decision-making across groups. For behavior, tVNS selectively improved decision accuracy in contexts characterized by initially low performance, without affecting response times. Drift diffusion modeling revealed that this improvement was specifically associated with increased drift rate, consistent with more efficient evidence accumulation with tVNS. These effects were consistent across groups but most pronounced when tVNS was applied at the early time point. Our results provide causal evidence that tVNS enhances decision-making in a state-dependent manner, likely by stabilizing attentional engagement and facilitating evidence accumulation when endogenous control is suboptimal.

## Introduction

The locus coeruleus (LC) is a small nucleus located in the brainstem which serves as the principal source of norepinephrine (NE) in the brain. Through its widespread projections to both cortical and subcortical regions (Aston-Jones and Cohen 2005, Breton-Provencher and Sur 2019), the LC-NE system serves as a core neuromodulatory pathway that regulates arousal and supports a wide range of behaviors (Aston-Jones, Rajkowski et al. 2000, Sara 2009, Sara and Bouret 2012, Tan, Adams et al. 2024). In recent years, growing evidence has implicated this system in perceptual decision-making, a cognitive domain that requires the accumulation of sensory evidence to decide on the most appropriate motor response (Hauser, Allen et al. 2017, van Kempen, Loughnane et al. 2019). However, the precise role of the LC-NE system in this context remains debated. A dominant view emerging from animal studies proposes that heightened LC-NE activity enhances the signal-to-noise ratio of sensory processing (Usher, Cohen et al. 1999, Clayton, Rajkowski et al. 2004, Ghosh and Maunsell 2022, Ghosh and Maunsell 2024), thereby improving decision accuracy, a mechanism known as “gain” modulation (Aston-Jones and Cohen 2005, Devilbiss 2019). In contrast, human studies have advanced an alternative “urgency” hypothesis, suggesting that increased LC-NE activity accelerates decision commitment and action release, potentially at the cost of accuracy (Murphy, Boonstra et al. 2016, Kelly, Corbett et al. 2021, O’Connell and Kelly 2021). These two accounts lead to divergent behavioral predictions, particularly regarding the impact of LC-NE activity on decision accuracy. Yet, direct empirical evidence in humans that can clearly distinguish between these competing views remains scarce.

One reason for the limited empirical support in humans for either hypothesis is that most studies have relied on correlational approaches, primarily using pupil size as an indirect index of LC-NE activity (Murphy, Boonstra et al. 2016, Ferrucci, Genovesio et al. 2021). While informative, such methods do not allow for definitive conclusions about the causal role of the LC-NE system. In recent years, however, transcutaneous vagus nerve stimulation (tVNS) has emerged as a promising non-invasive technique to causally probe LC function. This approach involves stimulation of the auricular branch of the vagus nerve at the level of the ear, which is known to enhance LC-NE activity via afferent projections to the nucleus tractus solitarii (NTS), which in turn projects to the LC (Yakunina, Kim et al. 2017, Skora, Marzecová et al. 2024, Pervaz, Thurn et al. 2025). Recently, using tVNS delivered as brief 4-second trains on each trial of a random dot motion task, we reported initial observations in humans that were consistent with the “gain” theory (Su, Vanvoorden et al., 2024). Specifically, online tVNS-induced increases in LC-NE activity were associated with improved decision accuracy without altering response speed. Notably, this improvement emerged only after errors, when accuracy in that condition had declined, effectively restoring performance to average levels, while having no measurable effect when performance was already adequate. This pattern aligns with the idea that LC-NE activity helps optimize the brain’s limited energetic resources by selectively enhancing performance when it benefits goal-directed behavior (Mather and Sutherland 2011, Mather, Clewett et al. 2016, Kaduk, Henry et al. 2023). However, while these initial findings provided first support for the gain hypothesis in humans, the selectivity of the effect could not be firmly characterized, and the study lacked sufficient power to rigorously examine the underlying computational mechanisms.

The present study involved a large sample of 62 healthy participants who performed the random dot motion task under tVNS and sham stimulation in separate blocks, with pupil size continuously recorded as a physiological proxy for LC-NE activity. Participants were instructed to respond as accurately and quickly as possible but received feedback only on accuracy, not speed, at the end of each trial. We hypothesized that tVNS would enhance decision accuracy specifically when performance is suboptimal, as indicated by higher error rates in the sham condition, and that our computational approach would reveal a selective effect on drift rate, reflecting enhanced evidence accumulation, consistent with the neural gain framework. Beyond this general prediction, we were further motivated by evidence that pupil size evolves dynamically throughout the course of goal-directed behavior (Murphy, Boonstra et al. 2016, Steinemann, O’Connell et al. 2018, Su, Vanvoorden et al. 2025), reflecting changes in LC-NE engagement over time. However, how transient neuromodulation by tVNS interacts with this evolving pattern of LC-NE engagement remains to be determined. To examine this, participants were assigned to one of three groups, each receiving tVNS or SHAM stimulation at a distinct time point during each trial: either well before, immediately before, or during decision-making. These timings corresponded to different stages of LC-NE engagement and allowed us to assess whether the timing of stimulation, and the associated pupil-linked LC-NE state, modulates tVNS effects on decision behavior and underlying computational mechanisms.

## Results

Sixty-two participants performed a random dot motion (RDM) task under tVNS and SHAM stimulation conditions (see Fig. 1A-B for more details). The RDM task required participants to discriminate the direction of coherently moving dots within a random display, as fast and accurately as possible. Reaction time (RT) and accuracy served as primary behavioral outcome measures, while pupil size was considered to assess LC-NE activity during the task in tVNS and SHAM blocks. Participants were assigned to one of three groups based on the timing of the 4-second tVNS/SHAM stimulation within each trial, corresponding to distinct phases of LC-NE engagement as informed by our previous study (Su, Vanvoorden et al., 2024): Group_Early_ (n = 20) received stimulation early in the trial, during the Fixation phase, when LC-NE activity was presumed to be at baseline. Group_Mid_ (n = 21), whose data were included in our previous study, received stimulation at a middle period spanning the Fixation and Pre-decision phases, aiming to target LC-NE as its activity began to rise. Group_Late_ (n = 21) received stimulation late in the trial, spanning Pre-decision and Decision phases, when LC-NE activity was reaching its task-related peak. Participants in each group received tVNS and SHAM stimulation in alternating blocks, allowing for within-subject comparisons in each group.

**Figure 1.**
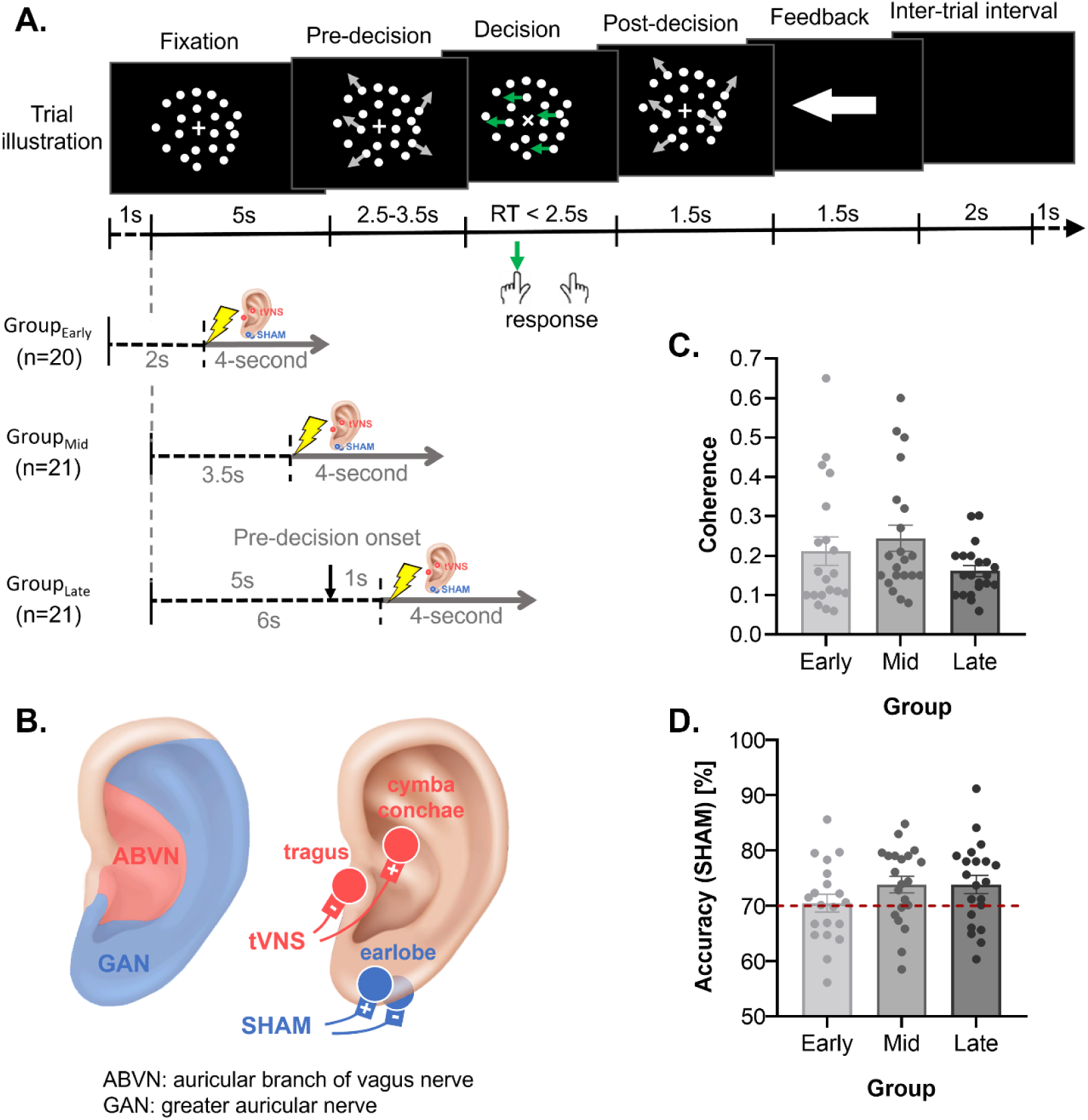
Experimental design. **A. Trial illustration in the random dot motion (RDM) task.** Participants were either part of Group_Early_ (n = 20), Group_Mid_ (n = 21), or Group_Late_ (n = 21), depending on the timing of tVNS/SHAM stimulation. Each trial began with a Fixation phase (lasting 6s in Group_Early_ and 5s in Group_Mid_ and Group_Late_), during which participants were instructed to focus on a centrally presented cross (+) surrounded by stationary dots. In the subsequent Pre-decision phase, the dots began moving randomly at a speed of 5 degrees per second, randomly drawn from a uniform distribution ranging from 2.5 to 3.5s. Then came the Decision phase, during which the central cross changed from the “+” to an “x”, and a proportion of dots began moving coherently in either a leftward or rightward direction (leftward in this example). The proportion of coherently moving dots in each trial was defined as coherence c’. Participants were required to identify the direction of coherent motion and respond by pressing a key with their left or right index finger (F12 or F5 on an inverted keyboard, respectively). They were instructed to respond as quickly and accurately as possible within a 2.5-second time limit to remain fully engaged in the task. After the response, the coherent motion transitioned back to purely random motion immediately after participants responded, initiating the Post-decision phase (1.5s). Feedback was then displayed for 1.5s in the form of a white arrow pointing left or right to indicate the actual motion direction for that trial. If participants failed to respond within the time limit, the feedback was replaced by the word “missed!”. The trial ended with an intertrial interval during which a black background was displayed (lasting 2s in Group_Early_ and 3s in Group_Mid_ and Group_Late_, to match overall trial duration in each group, despite the 1-second longer Fixation phase in Group_EARLY_). A 4-second train of tVNS/SHAM was applied during the task. In Group_Early_, participants received tVNS/SHAM stimulation early in the trial, 2s after the onset of the 6s Fixation phase. In Group_Mid_, participants received tVNS/SHAM stimulation at a middle time point, 3.5s after the 5s Fixation phase onset, thus covering the Fixation and Pre-decision phases. In Group_Late_, participants received stimulation late in the trial, 1s after the onset of the Pre-Decision phase, thereby spanning the Pre-decision and Decision phases. **B. tVNS setup.** Electrodes were put on the cymba conchae and tragus of the left ear in tVNS condition, allowing to generate an electric field covering the area innervated by the auricular branch of the vagus nerve. Electrodes were put on the left earlobe in the SHAM condition, covering the area of the greater auricular nerve. **C. Coherence in the RDM task.** Individual coherence data in Group_Early_, Group_Mid_, and Group_Late_ are represented by light, medium, and dark grey dots, respectively. Bar plots indicate the Mean ± SEM. Note comparable coherence values in the three groups. **D. Decision accuracy in SHAM blocks.** Note comparable decision accuracy of approximately 70% in SHAM blocks with calibrated coherence for the three groups.

To account for individual differences in visual perception, the coherence level (c’) of moving dots in the RDM task was adjusted at the beginning of the experiment for each participant to achieve approximately 70% baseline accuracy. This calibration procedure resulted in overall similar coherence levels across the three groups (One-way ANOVA, F(2, 59) = 2.090, p = 0.133, see Fig. 1C). Furthermore, as intended, accuracy in SHAM blocks averaged 73.86% with no significant differences across groups (One-way ANOVA, F(2, 59) = 2.681, p = 0.077, see Fig. 1D), despite slightly higher values in Group_Mid_ and Group_Late_ compared to Group_Early_.

### tVNS systematically increased pupil dilation through decision-making, regardless of its application time

Pupil size was continuously recorded throughout the RDM task. For all groups, pupil size was normalized within each participant using z-score and was then segmented into trials aligned either to the onset of stimulation or to the onset of coherent dot motion (Decision phase onset). Pupil dilation was quantified as a change from pupil baseline, which was always computed over a 1-second window starting 1s after Fixation onset (see Methods and Fig. 2A-C, left panel).

**Figure 2.**
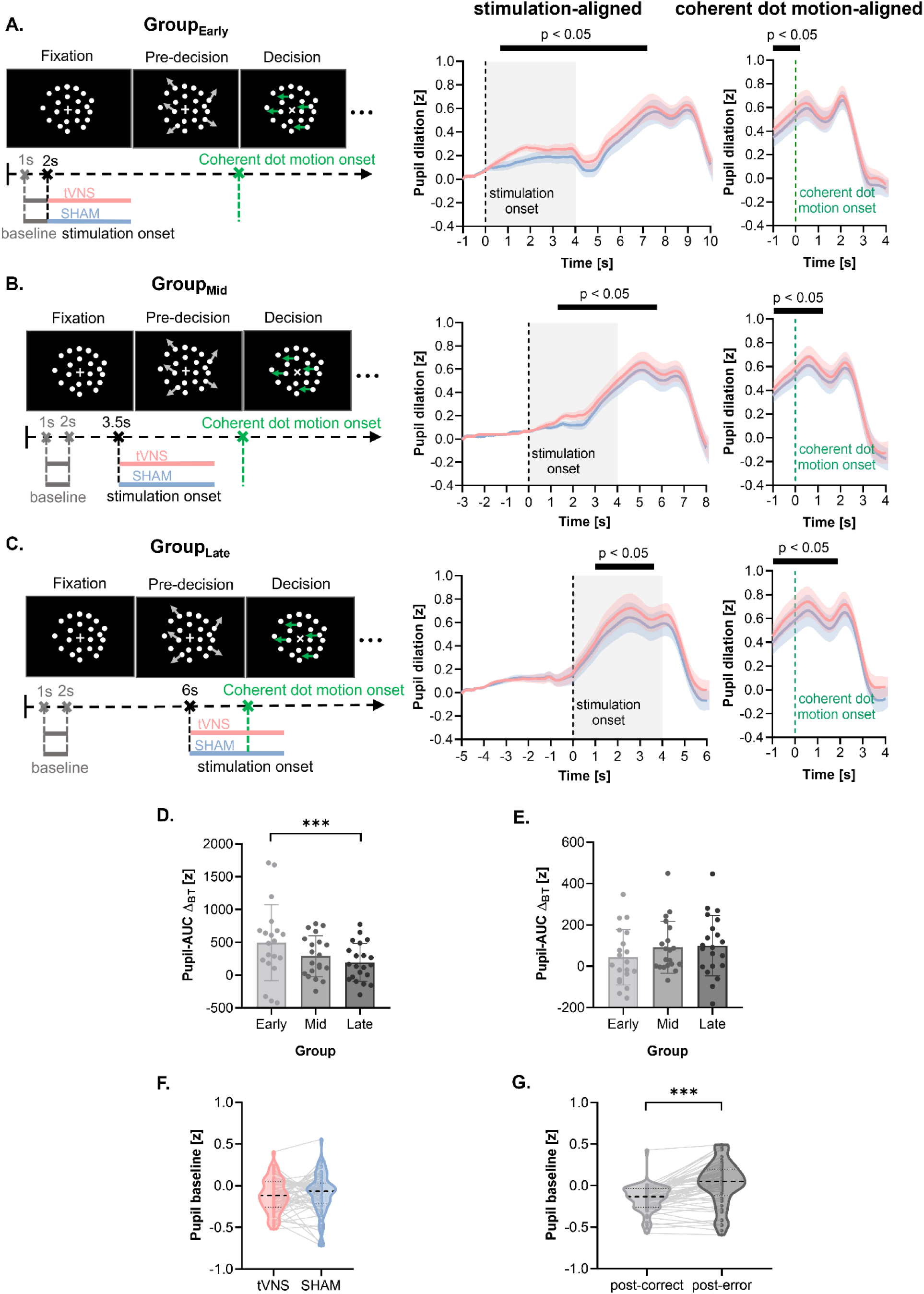
Pupil response in the RDM task. **A. Pupil dilation in Group_Early_.** The graphs display pupil dilation aligned to stimulation onset (middle panel) and coherent dot motion onset (right panel) in Group_Early_, separated for tVNS (red) and SHAM (blue) blocks. Shaded areas around the traces represent ± SEM. The gray transparent rectangle indicates the stimulation period. Black horizontal lines above the traces indicate the period showing significant differences between tVNS and SHAM blocks, identified using paired t-tests with cluster-based permutation correction (p < 0.05). Pupil dilation was significantly greater in tVNS than SHAM blocks from 0.62 s to 7.23 s after stimulation onset, and up to 0.16 s after motion onset. **B. Pupil dilation in Group_Mid_.** Same format as in A, for Group_Mid_. Significant tVNS-induced pupil dilation was observed from 1.21 s to 5.85 s post-stimulation onset, and up to 1.24 s post-coherent motion onset. **C. Pupil dilation in Group_Late_.** Same format as in A, for Group_Late_. Pupil dilation was significantly greater in tVNS than SHAM blocks from 0.98 s to 3.64 s after stimulation onset, and up to 1.85 s after coherent motion onset. **D. Pupil-AUC** Δ**_BT_ aligned to stimulation onset.** This graph shows the overall effect of tVNS on pupil dilation, expressed as the Pupil-AUC Δ_BT_, which corresponds to the area between the tVNS and SHAM curves, computed over the period showing significant tVNS effects in each group (indicated by the black horizontal lines above the traces in the middle panel of A-C). Each dot represents one participant from Group_Early_ (light gray), Group_Mid_ (medium gray), or Group_Late_ (dark gray); bars indicate group means ± SEM. Note the greater Pupil-AUC Δ_BT_ in Group_Early_ than Pupil-AUC Δ_BT_ than Group_Late_. **E. Pupil-AUC**Δ**_BT_ during decision-making.** Same format as in D, but here the Pupil-AUC Δ_BT_ was computed from the onset of coherent dot motion to the participant’s response. Note the comparable Pupil-AUC Δ_BT_ during decision-making across groups. **F. Block-Type effect on pupil baseline.** The graph displays median values as black dotted lines, distributions as violin plots, and individual data as thin gray lines. Note the comparable pupil baseline in tVNS (red) and SHAM (blue) blocks. **G. Trial-History effect on pupil baseline.** Same format as in F. Note the greater pupil baseline in post-error (dark gray) than post-correct trials (light gray). ***: 𝑝 < 0.001.

Pupil dilation consistently increased throughout the RDM task trials, peaking around 0.5s after coherent dot motion onset before gradually returning to baseline (see Fig. 2A-C and Table 1). This pattern was evident across all groups, regardless of whether the pupil data was aligned to stimulation onset (Fig. 2A-C, middle panel) or coherent dot motion onset (right panel). More importantly, all groups exhibited significant differences in pupil dilation between tVNS and SHAM conditions when pupil data were aligned to stimulation onset (Fig. 2A-C, middle panel), with paired t-tests using cluster-based permutations revealing significantly greater pupil dilation under tVNS compared to SHAM (p < 0.05), from 0.62s to 7.23s in Group_Early_, from 1.21s to 5.85s in Group_Mid_, and from 0.98s to 3.64s in Group_Late_. These results indicate that the tVNS effect on pupil dilation consistently emerged shortly after stimulation onset (between 0.62s and 1.21s) and systematically persisted until approximately the time of the participant’s response. Hence, its duration varied across groups, appearing longest in Group_Early_ and shortest in Group_Late_. To assess the resulting difference in the overall magnitude of the tVNS-induced pupil dilation across groups, we computed the cumulative difference in pupil dilation between tVNS and SHAM conditions, over the period in which a significant tVNS effect was observed. Specifically, we calculated the area under the curve (AUC) of pupil dilation traces aligned to stimulation onset for each subject, separately for each Block-Type (BT: tVNS and SHAM). The SHAM AUC was then subtracted from the tVNS AUC to derive a delta (Δ) measure reflecting tVNS-induced pupil dilation, referred to as Pupil-AUC Δ_BT_. The ANOVA run on this measure revealed a main effect of Group (F(2, 58) = 6.901, p = 0.002), indicating a greater Pupil-AUC Δ_BT_ with early compared to late tVNS timing (see Fig. 2D, Group_Early_ vs. Group_Late_: t = 3.681, 𝑝_𝑡𝑢𝑘𝑒𝑦_ = 0.001).

**Table 1.**
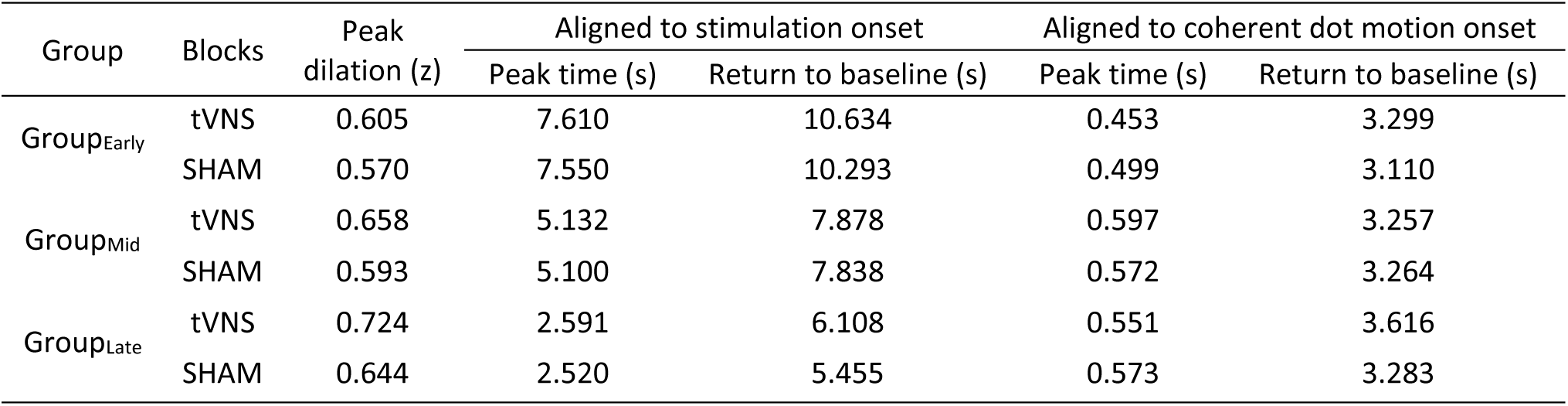
Summary of the dynamics of pupil dilation under tVNS and SHAM conditions across groups.

The tVNS effect was also evident in all groups when conducting paired t-tests with cluster-based permutations on pupil data aligned to coherent dot motion onset (Fig. 2A-C, right panel), with significant effects observed from -1s to 0.16s in Group_Early_, -1s to 1.24s in Group_Mid_, and -1s to 1.85s in Group_Late_ (all p < 0.05). Here too, we extracted a similar Pupil-AUC Δ_BT_ to assess the tVNS-induced pupil dilation specifically during the Decision phase, thus starting from the onset of coherent dot motion to the participants’ response. When restricting our analysis to this period, ANOVA revealed a comparable tVNS-related increase in pupil dilation across groups, as confirmed by the absence of Group effect on Pupil-AUC Δ_BT_ (F(2, 58) = 0.793, p = 0.458, Fig. 2E). Our results therefore provide strong evidence that, regardless of the timing of its application, the effect of tVNS tended to subside around the time of the response, even when applied early in the trial in Group_Early_. Hence, early stimulation (Group_Early_) led to a longer-lasting effect on pupil dilation such that the magnitude of the effect during decision-making was ultimately comparable across the three groups.

#### tVNS did not alter pupil baseline but the latter was increased following errors

Analyses on a subset of subjects in a prior study (Group_Mid_, n=20) suggested that using sufficiently long inter-stimulation intervals (∼ 11s) allows the tVNS effect on pupil to dissipate before the next trial begins (Su, Vanvoorden et al. 2025). In addition, this prior study reported a larger pupil baseline following errors compared to correct trials. To verify this in our larger dataset (n=62), we analyzed pupil baseline from all participants across groups, considering Block-Type (tVNS, SHAM) and Trial-History (post-correct, post-error) as within-subject factors. A linear mixed effects model confirmed the absence of a Block-Type effect on pupil baseline (F(1, 23.8) = 0.529, p = 0.474, see Fig. 2F). However, consistent with prior findings, we found a main effect of Trial-History, with higher pupil baseline following errors compared to correct trials (F(1, 17.3) = 86.666, p < 0.001, see Fig. 2G), irrespective of Block-Type (Block-Type × Trial-History interaction: F(1, 17069.4) = 1.457, p = 0.227).

### tVNS enhanced accuracy irrespective of its application time in the RDM task and without affecting RTs

To determine whether the activation of LC-NE by tVNS influences behavior and whether its effects depend on the timing of its application, accuracy and RTs were analyzed using generalized and linear mixed-effects models, respectively. Notably, accuracy was higher in tVNS blocks (Group_Early_ = 73.4 ± 1.5%, Group_Mid_ = 77.8 ± 1.8%, Group_Late_ = 76.0 ± 1.6%) compared to SHAM blocks (Group_Early_ = 70.5 ± 1.6%, Group_Mid_ = 75.4 ± 1.8%, Group_Late_ = 75.6 ± 1.9%) in all groups (see Fig. 3A), as confirmed by a significant main effect of Block-Type (χ^2^(1) = 8.77, p = 0.003) with no Block-Type × Group interaction (χ^2^ (2) = 1.145, p = 0.564) in the generalized mixed-effects model. Despite the relatively modest effect size, the improvement in accuracy under tVNS was consistent across participants in Group_Early_ (17/20) and Group_Mid_ (17/21), and still evident in a majority of Group_Late_ participants (13/21). In contrast, the linear mixed-effects model on RTs did not reveal any effect of Block-Type (F < 1.874, p > 0.154). As such, RTs in all groups were comparable in tVNS blocks (Group_Early_ = 1.105 ± 0.061s, Group_Mid_ = 1.314 ± 0.054s, Group_Late_ = 1.247 ± 0.045s) and SHAM blocks (Group_Early_ = 1.123 ± 0.057s, Group_Mid_ = 1.322 ± 0.049s, Group_Late_ = 1.238 ± 0.046s, Fig. 3B). However, we observed a main effect of Group, with participants in Group_Mid_ exhibiting the slowest overall RTs, while those in Group_Early_ responded the fastest (F(2, 15800.4) = 327.29, p < 0.001), regardless of the Block-Type (Block-Type × Group interaction: F(2, 13551) = 1.175, p = 0.309).

**Figure 3.**
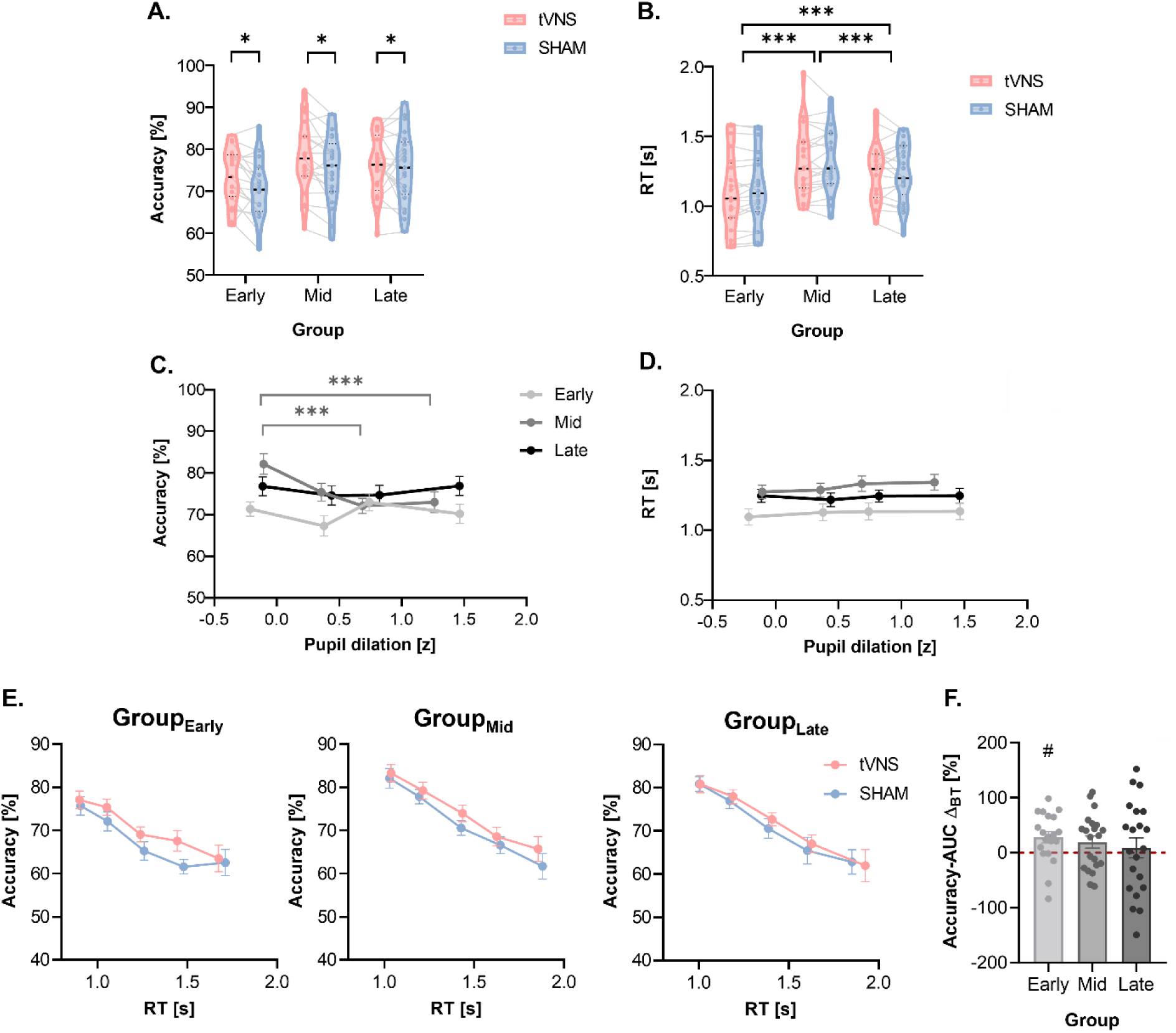
Accuracy and reaction times (RT) in the RDM task. **A. General tVNS effect on accuracy across groups.** The data features median values as black dotted lines, distributions as violin plots, and individual data as thin gray lines, separated for tVNS (red) and SHAM (blue) blocks in each group. Note the greater accuracy in tVNS compared to SHAM blocks in all groups. **B. General tVNS effect on RT across groups**. Same format as in A. Note comparable RTs in tVNS and SHAM blocks in all groups. **C. Effect of pupil dilation on accuracy in SHAM blocks.** Data are displayed as Mean ± SEM across pupil dilation bins for Group_Early_ (light gray), Group_Mid_ (medium gray), and Group_Late_ (dark gray) groups. Accuracy remained stable across pupil dilation bins in Group_Early_ and Groups_Late_, whereas a trend toward reduced accuracy with greater pupil dilation was observed in Group_Mid_. **D. Effect of pupil dilation on RT in SHAM blocks.** Same format as in C. RTs remained stable across pupil dilation bins in all groups. **E. tVNS effect on conditional accuracy curves across groups.** Accuracy is plotted as a function of RT, with each data point representing the mean ± SEM of accuracy within RT bins. In all groups, the conditional accuracy curve under tVNS (red) was shifted upward relative to SHAM (blue), indicating an improved speed-accuracy tradeoff with greater accuracy at comparable RTs. **F. Accuracy-AUC Δ_BT_ across groups.** Bars and individual dots represent the mean ± SEM of delta accuracy (tVNS minus SHAM) integrated across all RT bins. While there was no significant difference in Accuracy-AUC Δ_BT_ across groups, only Group_Early_ showed values significantly above zero. *: 𝑝 < 0.05. ***: 𝑝 < 0.001. #: 𝑝 < 0.05.

Following these findings, we checked whether the enhanced accuracy observed in tVNS blocks could result from the pupil being physically larger than in SHAM blocks during decision-making in the RDM task. In other words, did subjects respond more accurately with tVNS simply because their pupil was more dilated in these blocks? To control for this potential bias, we employed a generalized mixed-effects model to examine the relationship between accuracy and pupil dilation during decision-making in SHAM blocks. The analysis revealed a significant Pupil dilation × Group interaction on accuracy (χ^2^ (2) = 7.29, p = 0.026), indicating that, if anything, accuracy tended to decrease (rather than increase) with greater pupil size (Fig. 3C), an effect only significant in Group_Mid_ (bin1 VS. bin3: 𝑧 = 4.512, 𝑝_𝑡𝑢𝑘𝑒𝑦_ < 0.001; bin1 VS. bin4: 𝑧 = 4.046, 𝑝_𝑡𝑢𝑘𝑒𝑦_ = 0.003). A linear mixed-effects model did not reveal any effects of pupil dilation on RTs in SHAM blocks (F < 3.05, p > 0.07, Fig. 3D). Therefore, the behavioral effects observed in tVNS blocks are unlikely to result from physically larger pupils in this condition. Instead, tVNS appeared to enhance subjects’ decision accuracy without compromising RTs, as further illustrated by an upward shift in the conditional accuracy curve of the three groups (see Fig. 3E). To assess the overall impact of tVNS on decision accuracy across the RT distribution, we computed the difference in the area under the conditional accuracy curve (AUC) between each Block-Type (BT: tVNS and SHAM). Specifically, for each subject, we first calculated the delta of accuracy between tVNS and SHAM blocks within each RT bin, based on the average RTs obtained in the two types of blocks. These values were then integrated across the RT range using the trapezoidal rule to obtain a single summary measure, referred to as Accuracy-AUC Δ_BT_. This metric captures the overall magnitude of tVNS-induced accuracy changes over the temporal profile of decisions. A one-way ANOVA revealed no significant difference in Accuracy-AUC Δ_BT_ across groups (F(2, 59) = 0.487, p = 0.617, see Fig. 3F), suggesting that the overall effect of tVNS on shifting the conditional accuracy curve was comparable across groups with different tVNS timings. However, one-sample t-tests against zero indicated that only Group_Early_ showed a significantly positive Accuracy-AUC Δ_BT_ (t = 2.769, p = 0.012), while the effects in Group_Mid_ (t = 1.778, p = 0.091) and Group_Late_ (t = 1.602, p = 0.121) did not reach statistical significance.

#### tVNS improved accuracy in conditions with initially lower accuracy

The generalized mixed-effects model on accuracy also considered Trial-History (TH: post-correct, post-error) as a fixed effect and incorporated “Accuracy Δ_TH_” as a covariate. This covariate represents each participant’s overall delta of accuracy between trials following errors compared to trials following correct trials (post-error minus post-correct) in SHAM blocks (see Fig. 4A, blue dots) and was used to identify which condition (post-error or post-correct) yielded the lowest accuracy for each individual in the absence of tVNS. A negative Accuracy Δ_TH_ indicates lower accuracy in post-error compared to post-correct trials, whereas a positive Accuracy Δ_TH_ reflects the opposite pattern, with lower accuracy in post-correct trials. Notably, there was a significant Block-Type × Trial-History × Accuracy Δ_TH_ interaction (χ^2^ (1) = 50.316, p < 0.001); the quadruple interaction with Group was not significant (χ^2^ (2) = 1.311, p = 0.519). To explore this interaction, participants were separated into two fairly balanced clusters depending on the sign of their Accuracy Δ_TH_ values (see Fig. 4A): 32 participants showed negative values, indicating lower accuracy (LowACC) in post-error (PE) trials (LowACC-PE cluster), while 30 participants exhibited positive values, indicating lower accuracy in post-correct (PC) trials (LowACC-PC cluster). A comparison of coherence values (c’) showed a trend toward higher coherence (F(1, 60) = 3.693, p = 0.059) in the LowACC-PE cluster (0.237 ± 0.027) than in the LowACC-PC cluster (0.172 ± 0.019), consistent with our previous finding that higher coherence leads to poorer performance after errors in our task (Su, Vanvoorden et al. 2025). As expected by definition, within SHAM blocks, participants in the LowACC-PE cluster exhibited significantly lower accuracy in post-error compared to post-correct trials (𝑧 = 8.246, 𝑝_𝑡𝑢𝑘𝑒𝑦_ < 0.001, Fig. 4B, left panel), while participants in the LowACC-PC cluster showed lower accuracy in post-correct than in post-error trials (𝑧 = -3.480, 𝑝_𝑡𝑢𝑘𝑒𝑦_ = 0.003, Fig. 4B, right panel). More interestingly, tVNS selectively enhanced accuracy in post-error trials of LowACC-PE participants (post-correct: 𝑧 = -2.018, 𝑝_𝑡𝑢𝑘𝑒𝑦_ = 0.181; post-error: 𝑧 = 4.687, 𝑝_𝑡𝑢𝑘𝑒𝑦_ < 0.001), while it selectively improved accuracy in post-correct trials of LowACC-PC participants (post-correct: 𝑧 = 4.055, 𝑝_𝑡𝑢𝑘𝑒𝑦_ < 0.001; post-error: 𝑧 = -1.053, 𝑝_𝑡𝑢𝑘𝑒𝑦_ = 0.718). Hence, the tVNS train enhanced accuracy in both participant clusters, but did so in different conditions, specifically in the one where they initially showed lower accuracy in SHAM blocks (LowACC trials, surrounded by red rectangles in Fig. 4B). Consequently, in tVNS blocks, the Accuracy Δ _TH_ became less negative for LowACC-PE participants and less positive for LowACC-PC participants (see Fig. 4A, red dots).

**Figure 4.**
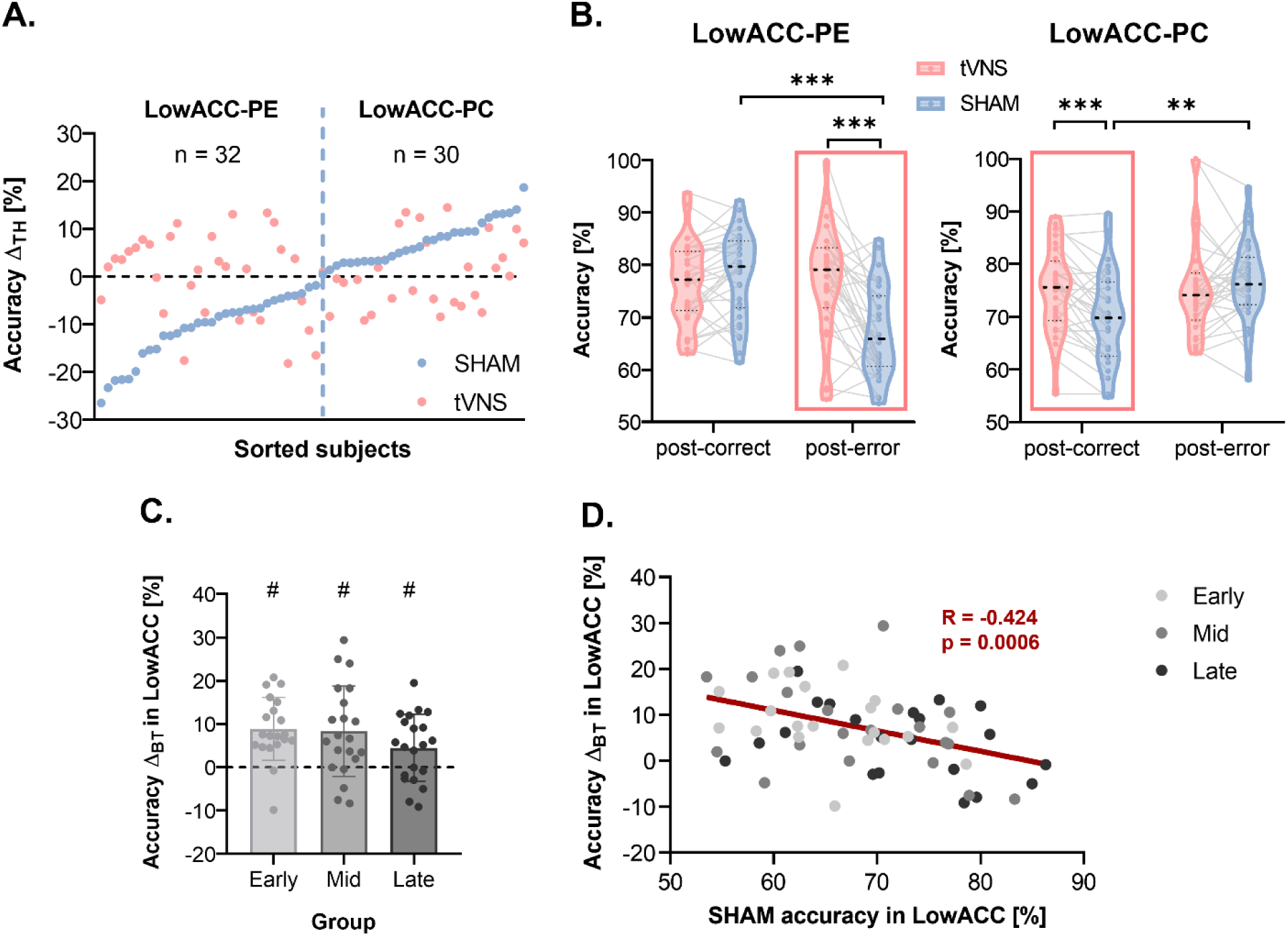
Accuracy as a function of Trial-History (TH) in the RDM task. **A. Definition of Participant-Cluster based on Accuracy Δ_TH_ values.** Dots represent Accuracy Δ_TH_ values in each participant, separated for tVNS (red) and SHAM (blue) blocks. Accuracy Δ_TH_ reflects the delta (Δ) of accuracy between post-error and post-correct trials (post-error minus post-correct). Participants were sorted and then clustered based on the sign of their Accuracy Δ_TH_ values in SHAM blocks. A negative value (left, n = 32) indicates lower accuracy in post error trials (LowACC-PE cluster), while a positive value (right, n = 30) indicates lower accuracy in post correct trials (LowACC-PC cluster). **B. Effect of tVNS on accuracy in each Participant-Cluster, as a function of Trial-History.** The accuracy data (in %) features median values as black dotted lines, distributions as violin plots, and individual data as thin gray lines, separated for tVNS (red) and SHAM (blue) blocks in each group. Note that tVNS selectively improved accuracy in post-error trials for LowACC-PE participants (marked with a red rectangle in the left panel), while selectively improved accuracy in post-correct trials for LowACC-PC participants (marked with a red rectangle in the right panel). **C. Accuracy Δ_BT_ in LowACC trials across groups.** Accuracy Δ_BT_ represents the delta of accuracy between tVNS and SHAM blocks (tVNS minus SHAM). Data were specifically derived from LowACC trials (post-error trials in LowACC-PE participants and post-correct trials in LowACC-PC participants). Individual data in Group_Early_, Group_Mid_, and Group_Late_ are represented by light, medium, and dark gray dots, respectively. Bar plots indicate the Mean ± SEM. Note that Accuracy Δ_BT_ values were comparable across groups, and each group showed significant values over 0. **D. Correlation between SHAM accuracy and Accuracy Δ_BT_ in LowACC trials.** Note that participants who exhibited the lower accuracy in SHAM blocks were those who benefited the most from the tVNS, as reflected by the greater Accuracy Δ_BT_ in these subjects. **: 𝑝 < 0.01. ***: 𝑝 < 0.001. #: 𝑝 < 0.05.

Following this finding, we restricted the next analyses to the trial type showing lower accuracy in SHAM blocks (LowACC trials) for each participant. That is, we selected the post-error trials in participants of the LowACC-PE cluster and the post-correct trials in participants of the LowACC-PC cluster and computed the Accuracy Δ_BT_, defined as the difference in accuracy between Block-Type (BT: tVNS−SHAM) for these LowACC trials. The value of Accuracy Δ_BT_ reflects the degree to which tVNS enhanced accuracy in each participant. We first compared these values between the three different groups (Early, Mid, Late), but the one-way ANOVA did not reveal any significant difference (F(2, 59) = 1.598, p = 0.211, see Fig. 4C). In fact, t-tests against zero showed that the Accuracy Δ_BT_ was significant in each group (all t > 2.664, all p < 0.045). Hence, tVNS rescued accuracy in LowACC trials, whatever the timing at which it was applied. Finally, in the last analysis, we pooled all participants from the three groups together and ran a Pearson’s partial correlation analysis to investigate the degree to which the Accuracy Δ_BT_ in LowACC trials depended on how low the accuracy was in these trials during SHAM blocks. Interestingly, participants who exhibited the lower accuracy in SHAM blocks were those who benefited the most from the tVNS, as reflected by the greater Accuracy Δ_BT_ in these subjects (R = -0.424, p = 0.0006, Fig. 4D). Hence, tVNS helped participants in low accuracy trials, especially in those showing lowest accuracy, regardless of the timing at which it was applied.

The Block-Type × Trial-History × Accuracy Δ_TH_ interaction was not significant for RTs (F(1, 17423) = 0.294, p = 0.587), but the linear mixed-effects model revealed a significant Group × Trial-History interaction (F(2, 11482.7) = 5.582, p = 0.004). Overall, participants displayed longer RTs following errors compared to correct trials, consistent with the “post-error slowing” phenomenon repeatedly reported in past research (Adkins, Zhang et al. 2024, Dyson 2024, Rafiezadeh, Tashk et al. 2024). This post-error slowing effect reached significance in Group_Mid_ and Group_Late_ (Group_Mid_: t(48.8) = -3.954, 𝑝_𝑡𝑢𝑘𝑒𝑦_ = 0.003; Group_Late_: t(43.6) = 5.341, 𝑝_𝑡𝑢𝑘𝑒𝑦_< 0.001; see middle and right panels in Fig. 5). In Group_Early_, although a similar trend was visually present (post-correct: 1.103 ± 0.059s, post-error: 1.144 ± 0.062s), the effect was not statistically significant (t(42.5) = -2.243, 𝑝_𝑡𝑢𝑘𝑒𝑦_ = 0.24, see left panel in Fig. 5). Importantly, none of these effects interacted with the Block-Type (F < 2.203, p > 0.111), confirming the absence of tVNS effect on RTs (see Fig. 3B). In conclusion, tVNS selectively enhanced decision accuracy without incurring a cost on RT, as also reflected by the tVNS-related upward shift of the conditional accuracy curve when specifically considering LowACC trials in all experimental groups (see Fig. 6A). We further analyzed the Accuracy-AUC Δ_BT_ (tVNS minus SHAM) derived from these conditional accuracy curves involving only LowACC trials. A one-way ANOVA revealed a significant group effect on Accuracy-AUC Δ_BT_ (F(2, 59) = 3.467, p = 0.038, Fig. 6B), with the Group_Early_ exhibiting a significantly larger delta compared to the Group_Late_ (t = 2.601, 𝑝_𝑡𝑢𝑘𝑒𝑦_= 0.031). One-sample t-tests against zero further confirmed that both Group_Early_ (t = 5.743, p < 0.001) and Group_Mid_ (t = 3.857, p < 0.001) exhibited significantly positive Accuracy-AUC Δ_BT_ values, while the effect in Group_Late_ did not reach significance (t = 1.895, p = 0.073). These results suggest that when only considering LowACC trials, the magnitude of tVNS-induced accuracy enhancement across the RT range was more pronounced when stimulation was applied earlier during the decision process.

**Figure 5.**
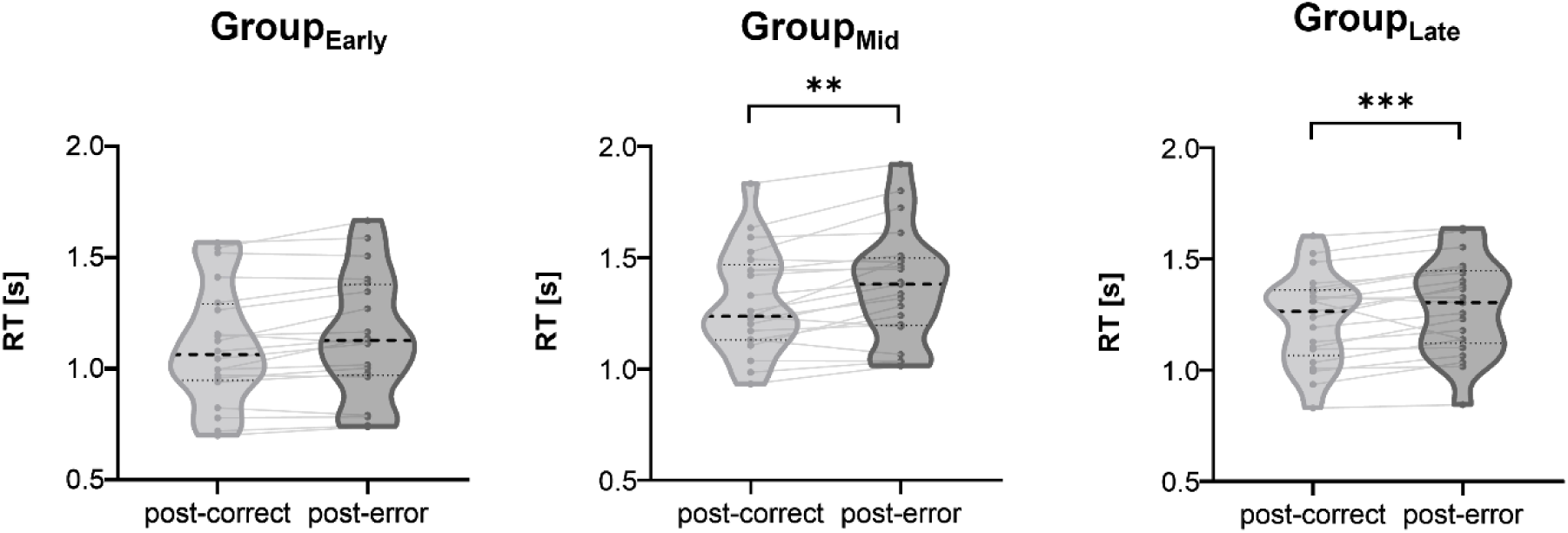
Reaction time (RT) as a function of Trial-History in the RDM task. The RT data features median values as black dotted lines, distributions as violin plots, and individual data as thin gray lines. Note significantly longer RTs in post-error (dark gray) compared to post-correct (light gray) trials in Group_Mid_ (middle panel), and Group_Late_ (right panel), while participants in Group_Early_ trended to make slower responses without significance (left panel). **: 𝑝 < 0.01. ***: 𝑝 < 0.001.

**Figure 6.**
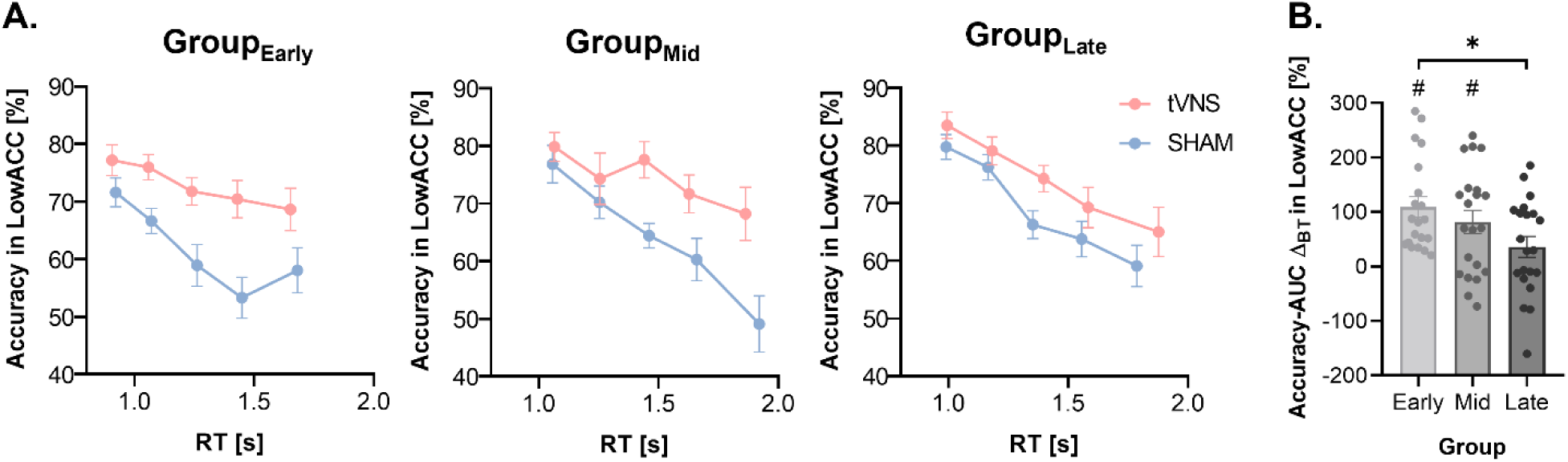
tVNS effect on accuracy across RT spectrum. **A. Conditional accuracy curves in LowACC trials.** Data are derived from LowACC trials in Group_Early_ (left panel), Group_Mid_ (middle panel), and Group_Late_ (right panel), with separate curves for tVNS (red) and SHAM (blue) blocks. Each data point represents the Mean ± SEM of accuracy calculated within RT bins. Note that the conditional accuracy curve under tVNS shifts upward relative to SHAM in all groups. **B. Accuracy-AUC** Δ**_BT_ in LowACC trials across groups.** Bars represent the Accuracy-AUC Δ_BT_ as Mean ± SEM, while dots are used to display individual data, with values corresponding to the delta of conditional accuracy curves (tVNS minus SHAM) in LowACC trials integrated across RT bins. Note that Accuracy-AUC Δ_BT_ values were significantly larger than 0 in Group_Early_ and Group_Mid_, and that Group_Early_ exhibited significantly larger Accuracy-AUC Δ_BT_ values compared to Group_Late_. *: 𝑝 < 0.05. #: 𝑝 < 0.05.

#### tVNS improved drift rate in conditions with initially lower accuracy and drift rate, regardless of its application time

We exploited the drift diffusion model (DDM) to explore the computational mechanisms underlying the selective behavioral effects. The DDM provides three key parameters: drift rate (reflecting the efficiency of sensory evidence accumulation) (Ratcliff and McKoon 2008), boundary (representing decision-making criteria) (Ratcliff, Smith et al. 2016), and non-decision time (capturing processes other than decision-making, like motor execution) (Hoxha, Chevallier et al. 2023). Here, we used a refined DDM with linearly collapsing boundaries, where a lower boundary intercept can be interpreted as a higher initial urgency, and a steeper boundary slope can be seen as a stronger increase in urgency to respond as time progresses, leading to a less skewed RT distribution and a drop in accuracy over time (Ratcliff, Smith et al. 2016). A two-way repeated measures ANOVA on drift rate, with Block-Type (tVNS, SHAM) and Trial-Type (LowACC, HighACC) as within-subject factors, revealed significant main effects (Block-Type: F(1, 48) = 7.733, p = 0.008; Trial-Type: F(1, 48) = 18.960, p < 0.001) and a Block-Type × Trial-Type interaction (F(1, 48) = 28.883, p < 0.001). Post-hoc comparisons showed that drift rate was generally lower in LowACC compared to HighACC trials for SHAM blocks (t = -6.454, 𝑝_ℎ𝑜𝑙𝑚_ < 0.001, see Fig. 7A), Crucially, tVNS significantly increased drift rate relative to SHAM in LowACC trials (t = 5,436, 𝑝_ℎ𝑜𝑙𝑚_ < 0.001), effectively restoring it to levels comparable to HighACC trials in tVNS blocks (t = 1.590, 𝑝_ℎ𝑜𝑙𝑚_ = 0.237). In contrast, no significant difference in drift rate was observed between tVNS and SHAM in HighACC trials (t = -2.100, 𝑝_ℎ𝑜𝑙𝑚_ = 0.123), suggesting that the tVNS effect was specific to conditions in which baseline performance was suboptimal. For the remaining model parameters, including boundary intercept, boundary slope, and non-decision time, no significant main effects or interactions were found (all F < 0.047, all p > 0.828; Fig. 7B-D).

**Figure 7.**
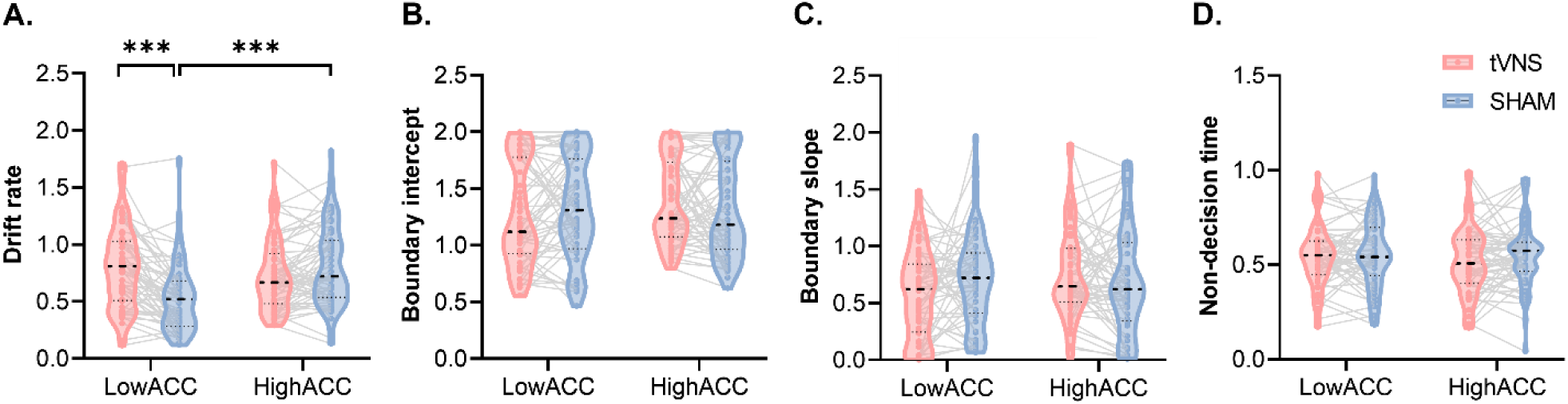
Drift Diffusion Model (DDM) parameters. The DDM data features median values as black dotted lines, distributions as violin plots, and individual data as thin gray lines. **A. Drift rate.** Note the lower drift rate in LowACC than HighACC trials for SHAM blocks, and the selective tVNS increase in drift rate in the LowACC trials only. **B. Boundary intercept.** Note the comparable values between tVNS and SHAM blocks in LowACC and HighACC trials**. C. Boundary slope.** Same result as in B. **D. Non-decision time.** Same result as in B. ***: 𝑝 < 0.001.

#### tVNS induced a similar pupil dilation in trial types with low or high accuracy

Given the selective effect of tVNS on behavior, we asked whether its impact on pupil-linked LC-NE activity was similarly selective. To do so, we first analyzed Pupil-AUC Δ_BT_ measures as a function of Trial-History and Group, and by considering the individual Accuracy Δ_TH_ in SHAM blocks (see Fig. 4A) as covariable. Repeated-measures ANOVAs did not reveal any significant effect involving Trial-History or Accuracy Δ_TH_ on Pupil-AUC Δ_BT_, whether considering data aligned to stimulation onset or during the Decision phase (all ANOVA F < 1.895, all p > 0.174), suggesting that pupil responses to tVNS were not modulated by the outcome of the preceding trial and how it affected accuracy. To ascertain this point, we also tested whether tVNS elicited larger pupil responses in trials characterized by initially lower accuracy (LowACC: post-error trials for participants in LowACC-PE cluster and post-correct trials for participants of LowACC-PC cluster) compared to those with initially higher accuracy (HighACC: post-correct trials for participants in LowACC-PE cluster and post-error trials for participants in LowACC-PC cluster). As such, we compared Pupil-AUC Δ_BT_ between LowACC and HighACC trials across groups, conducting two separate repeated-measures ANOVAs for stimulation-aligned (Fig. 2D) and Decision phase (Fig. 2E) measures, with Trial-Type (LowACC, HighACC) and Group (Early, Mid, Late) as factors. Apart from a significant main effect of Group on stimulation-aligned Pupil-AUC Δ_BT_ (F(2, 58) = 6.901, p = 0.002, Fig. 8A), no significant main effects or interactions involving Trial-Type were detected in either analysis (all F < 1.252, all p > 0.268; see Fig. 8A– B). These results suggest that tVNS induced similar pupil dilations in trials with initially low or high accuracy, despite selectively enhancing decision accuracy only in the former. Thus, the selective tVNS-related gain in accuracy was not accompanied by a corresponding selective increase in pupil responses. Instead, tVNS seems to have triggered a consistent increase in LC-NE activity across trial types.

**Figure 8.**
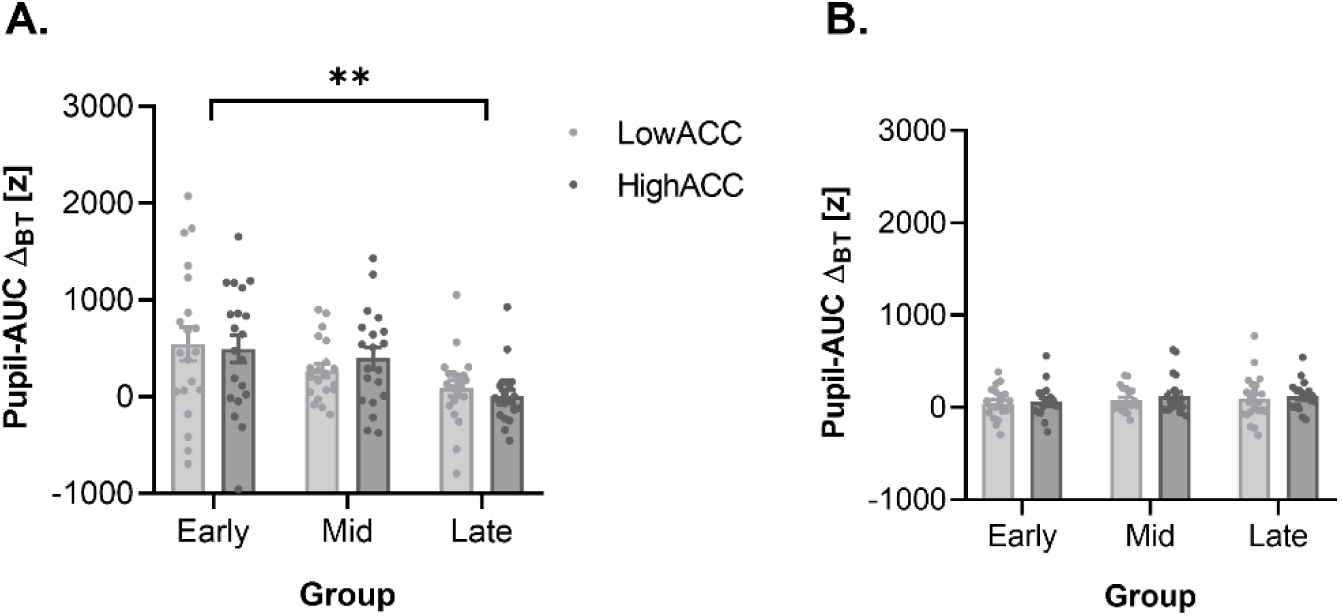
tVNS effect on pupil dilation across trial types and experimental groups. These graphs show the overall effect of tVNS on pupil dilation, expressed as the Pupil-AUC Δ_BT_. Individual dots and bars represent values for LowACC trials (light gray; post-error trials in LowACC-PE participants and post-correct trials in LowACC-PC participants) and HighACC trials (dark gray; post-correct trials in LowACC-PE participants and post-error trials in LowACC-PC participants) across Group_Early_, Group_Mid_, and Group_Late_. Bar plots indicate Mean ± SEM. **A. Pupil-AUC** Δ**_BT_ aligned to stimulation onset.** While a significantly higher Pupil-AUC Δ_BT_ was observed in Group_Early_ when compared to Group_Late_, all groups showed comparable Pupil-AUC Δ_BT_ between LowACC and HighACC trials. **B. Pupil-AUC** Δ**_BT_ during the Decision phase.** All groups showed comparable Pupil-AUC Δ_BT_ between LowACC and HighACC trials. **: *p* < 0.01.

## Materials and methods

### Participants

Sixty-two healthy participants were considered in this study (37 women, 24.8 ± 0.78 years old). Data from 21 of these participants were already exploited in a recent study (Su, Vanvoorden et al. 2025). All participants reported normal or corrected-to-normal vision, no history of neurological or psychiatric disorders, and no physical impairments. Before the experiment, they provided written informed consent and received financial compensation for their participation. They were all naive to the purpose of the study. The experimental protocol was approved by the Ethics Committee of the Université Catholique de Louvain (UCLouvain), Brussels, Belgium (approval numbers: 2018/22MAl/219 and 2023/13JUL/322), in accordance with the Declaration of Helsinki.

### Experimental protocol

#### Experimental setup

Experiments were conducted in a quiet and dimly lit room. The participants were seated at a table facing a 23-inch computer screen (60Hz refresh rate) positioned 60cm from their eyes, which displayed the visual stimuli of the random dot motion (RDM) task (see Fig. 1A). They rested their forearms on the table, palms facing down, with the response keys F12 and F5 positioned under their left and right index fingers, respectively. During the task, tVNS/SHAM stimulation was delivered and pupillometry was continuously recorded, participants were instructed to minimize any movements and eye blinks during the recordings.

#### Transcutaneous vagus nerve stimulation

Pulses were generated using a Digitimer (Digitimer Ltd., model DS3), triggered by a Master8 device (A.M.P.I., model Master-8). As shown in Figure 1B, in tVNS condition, the electrodes were placed at the left cymba conchae (anode) and tragus (cathode), which are heavily innervated by the auricular branch of the vagus nerve (Safi, Ellrich et al. 2016, Badran, Brown et al. 2018, Kreisberg, Esmaeilpour et al. 2021). In SHAM condition, the electrodes were placed at the left earlobe, which is not expected to induce brainstem or cortical activation (Sellaro, van Leusden et al. 2015, Steenbergen, Sellaro et al. 2015, Sharon, Fahoum et al. 2021). Pulses (width, 200𝜇s) were delivered at a rate of 25Hz for 4s. To ensure similar sensory perception between tVNS and SHAM conditions, stimulation intensity was calibrated separately for each subject and condition, as also reported in (Su, Vanvoorden et al. 2024). Specifically, participants underwent a "method of limits" procedure to determine the tVNS intensity at a level experienced as just below painful (Yakunina, Kim et al. 2017, Ventura-Bort, Wirkner et al. 2018). Stimulation began at 0.1 mA and increased in 0.2 mA steps until participants reported a sensation of 9 on a 10-point scale (0 = no sensation, 10 = painful). This intensity was repeated before a decreasing series, where stimulation was lowered in 0.2 mA steps until the sensation dropped to 6 or below. The final tVNS intensity was set as the average of the intensities rated as 8. For SHAM intensity, the same procedure was used, but participants described the sensation relative to tVNS without providing numerical ratings. The final SHAM intensity was set as the average of the intensities they perceived as similar to tVNS at 8.

#### Random dot motion task

The RDM task was implemented by means of Matlab7 (The Mathworks Inc. Natick, Massachusetts, USA) and the Psychophysics toolbox (Brainard and Vision 1997). As shown in Figure 1A, each trial began with a Fixation phase, during which participants fixated a central cross surrounded by stationary dots. This was followed by a Pre-decision phase, where dots moved randomly at 5 degrees per second. Then, during the Decision phase, a proportion of dots began moving coherently leftward or rightward, and participants were required to discriminate the coherent direction and provide their response by pressing the response key (left: F12, right: F5) as quickly and accurately as possible within 2.5s. The difficulty of these discriminations was set by the proportion of coherently moving dots (coherence c’) which was adjusted individually (see below). After responding, the motion became random again during a Post-decision phase, followed by a Feedback phase displaying the correct direction or "missed!" in the absence of response. Finally, an inter-trial interval consisting of a black screen was presented to separate trials. A 4-second tVNS/SHAM stimulation was applied in each trial, with participants randomly assigned to one of three groups based on the timing of this train within the trial. Stimulation began 2s, 3.5s, and 6s after trial onset for Group_Early_, Group_Mid_, and Group_Late_, respectively. Consequently, Group_Early_ (n = 20, 14 women; 23.4 ± 0.60 years old) received stimulation exclusively during the Fixation phase, Group_Mid_ (n = 21, 12 women; 25.7 ± 0.99 years old) during both the Fixation and Pre-decision phase, and Group_Late_ (n = 21, 11 women; 25.4 ± 0.75 years old) during the Pre-decision and Decision phases. Participants in each group received tVNS and SHAM stimulation in alternating blocks, allowing for within-subject comparisons in each group. The data from Group_Mid_ were previously exploited in an earlier study (Su, Vanvoorden et al. 2024). Note that the task was the exact same in all groups but for the duration of the Fixation that was 1s longer in the Group_Early_ (6s instead of 5s in the two other groups) to allow for the tVNS train to occur fully during this phase, while still having a baseline period from 1-2s after Fixation onset for pupil analyses. Yet, the overall trial duration was matched between groups by having a 1s shorter intertrial interval in Group_Early_ (see below).

#### Experimental design

All participants came to the lab for a session of about 3 hours, which always started with some initial practice and difficulty calibration routines. The practice consisted of 4 blocks of 40 trials, with coherence levels (c’) progressively decreasing from 0.8 to 0.1 (0.8, 0.4, 0.2, 0.1) to familiarize participants with the task. Then, to account for individual differences in visual perception, the coherence level (c’) of moving dots in the RDM task was adjusted at the beginning of the experiment for each participant to achieve approximately 70% baseline accuracy. This was done in two calibration blocks of 100 trials, where coherence levels (c’ = 0.1, 0.2, 0.4, 0.6, 0.8) were presented in equal proportions and randomized across trials. A proportional rate diffusion model was then fitted to the data from these calibration blocks to estimate each participant’s psychometric function (Palmer, Huk et al. 2005), from which the c’ value yielding 70% accuracy was interpolated and used in the main experiment. Following difficulty calibration, participants underwent a stimulation intensity calibration, where individual tVNS and SHAM intensities were determined to ensure consistent sensory perception across conditions (as described above). During the main experiment, participants completed the RDM task in 8 blocks of 40 trials, with 4 blocks of tVNS and 4 blocks of SHAM stimulation presented in an alternating sequence. To control for order effects, the sequence of the first 4 blocks was reversed in the second half of the session, and block order was counterbalanced across participants. Short breaks were provided between blocks to minimize fatigue.

### Behavioral measures and analyses

RT and accuracy in the RDM task were assessed as primary behavioral measures. Trials with missing responses or excessively fast reactions (< 450 ms) were excluded from both behavioral and pupillometric analyses. RT was defined as the time between the onset of coherent motion and the participant’s key press, while accuracy was calculated as the percentage of correct responses across valid trials. On average, 142.3 ± 3.38 trials in the tVNS condition and 141.9 ± 3.41 trials in the SHAM condition were available for behavioral analyses across all participants.

Trials were categorized by Block-Type (BT: tVNS, SHAM) and were further classified based on whether they followed a correct choice or an error in the preceding trial (Trial-History, TH: post-correct, post-error). Moreover, we computed “Accuracy Δ_TH_”, reflecting each participant’s overall difference in accuracy (delta, Δ) between Trial-History (post-error minus post-correct) in SHAM blocks (see Fig. 4A, blue dots). The Accuracy Δ_TH_ value was used as a covariate in our analyses; it also allowed us to ultimately categorize the participants in two clusters depending on whether they initially displayed a negative value, hence lower accuracy (LowACC) in post-error (PE) trials (n=32, LowACC-PE) or a positive value, hence lower accuracy in post-correct (PC) trials (n = 30, LowACC-PC).

Statistical analyses of behavioral data were conducted in Jamovi (version 2.5.3) using generalized (Accuracy) and linear (RT) mixed-effects models. These models included Block-Type (tVNS, SHAM), Group (Early, Mid, Late), and Trial-History (post-correct, post-error) as fixed factors, with individual Accuracy Δ_TH_ in SHAM blocks as a covariate. Post-hoc tests with Tukey corrections were applied to explore significant interactions, and the Participant-Cluster (LowACC-PE, LowACC-PC) was incorporated when necessary. Follow-up ANOVAs (and Bonferroni-corrected t-tests) were also run on Accuracy Δ_BT_ measures computed in each participant, defined as the difference in accuracy between Block-Type (tVNS minus SHAM). Moreover, relationships between variables were tested using Pearson’s partial correlation analyses.

To capture how tVNS influenced accuracy as a function of RT, we computed conditional accuracy curves. These conditional accuracy curves were generated separately for tVNS and SHAM blocks using a sliding-window approach. That is, for each participant, trials were sorted by RT within each block type, and mean accuracy was computed within overlapping windows spanning 40% of the RT range, shifting in 15% increments. This procedure yielded a total of 5 bins per participant and condition. The resulting conditional accuracy curves for tVNS and SHAM blocks were first computed at the individual level and then averaged across participants. SEM was calculated on accuracy values across participants for each bin. To quantify the overall effect of tVNS on these curves, we calculated the area under the accuracy curve (AUC) between Block-Type (tVNS minus SHAM) using trapezoidal integration over RT bin centers averaged across tVNS and SHAM conditions, referred to as Accuracy-AUC Δ_BT_. Accuracy-AUC Δ_BT_ values were compared across groups using a one-way ANOVA, with Tukey’s test used for post hoc comparisons. Results are expressed as means ± SEM throughout the manuscript, except for violin plots where medians and interquartile ranges (IQR) are shown. An alpha level of 0.05 was used for all statistical tests.

### Drift diffusion modeling

The DDM was fitted to the RT and Accuracy data using a variant that includes linearly collapsing decision boundaries. The decision boundary during each trial was modeled as:

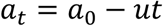

where 𝑎_𝑡_ is the boundary height at time 𝑡, 𝑎_0_ is the initial decision boundary (further referred to as boundary intercept), and 𝑢 ≥ 0 is the collapse rate of the boundary (further referred to as boundary slope). The drift rate, boundary intercept, boundary slope, and non-decision time were estimated separately for each participant within each condition defined by Block-Type (tVNS, SHAM) and Trial-History (post-correct, post-error). Quantile optimization was used for fitting the DDM (Ratcliff and Tuerlinckx 2002). The optimization was performed by a differential evolution algorithm as implemented in the DEoptim package in R (Mullen, Ardia et al. 2011). The population size was set at ten times the number of free parameters. Convergence was met if no improvement in the objective function was found in the last 100 generations. To ensure a reliable interpretation of estimated parameters, a parameter recovery analysis was performed. We simulated the data of 1000 simulated participants using parameter values randomly sampled from their typical range. We managed to recover the parameters with as few as 25 trials: all correlation coefficients between generated and estimated parameters were larger than 0.73, except for the boundary slope (r = 0.53). As a result, we excluded 10 subjects with fewer than 25 post-error trials, leaving 52 subjects with an average of 37.4 ± 1.41 and 40.1 ± 1.57 post-error trials in the tVNS and SHAM conditions, respectively. For post-correct trials, all 62 subjects were included, with an average of 107.9 ± 3.03 trials in the tVNS condition and 105.2 ± 3.07 trials in the SHAM condition. Two way repeated-measures ANOVAs were conducted to compare estimated DDM parameters across Block-Type (tVNS, SHAM), and Trial-Type (LowACC trials: post-error trials in the LowACC-PE cluster and post-correct trials in the LowACC-PC cluster, HighACC trials: post-correct trials in the LowACC-PE cluster and post-error trials in the LowACC-PC cluster).

### Pupillometry data acquisition and analyses

Pupil size was recorded binocularly at 1000 Hz using an Eyelink 1000+ eye-tracker (SR Research) and analyzed in MATLAB. Blink periods were identified and removed by linearly interpolating values 100ms before and after each blink, as implemented in custom MATLAB scripts. The pupil signal was then low-pass filtered using a 10 Hz fourth-order Butterworth filter with zero-phase shift. To ensure data quality, trials where interpolated data exceeded 50% of the total samples were excluded. For subsequent analyses, the eye with the more stable pupil signal, determined by the lower proportion of interpolated data and smaller deviation, was selected. After these pre-processing procedures, one participant in Group_Mid_ was excluded (remaining n = 20 in Group_Mid_ for pupil analyses). Additionally, after excluding trials with no response or very fast responses, an individual average of 140.7 ± 3.49 in tVNS blocks and 139.7 ± 3.51 trials in SHAM blocks were included for pupil analyses. To avoid arbitrary units, we converted pupil size into z scores computed across all conditions within each subject. Pupil baseline was defined as the average pupil size from 1-2s after trial onset in the three groups (see Fig. 2A-C, left panel). Then, to assess how tVNS influenced LC-NE activity during the RDM task, dynamics of pupil dilation in tVNS and SHAM blocks were analyzed in the three groups by considering group-specific 11-second windows aligned to stimulation onset: [-1 10]s for Group_Early_, [-3 8]s for Group_Mid_, and [-5 6]s for Group_Late_ (see Fig. 2A-C, middle panel). Moreover, to specifically consider pupil dilation during decision-making in all groups, we also used a 5-second window aligned to coherent dot motion onset ([-1 4]s in all groups, see Fig. 2A-C,right panel).

For the statistical analysis of pupil baseline, we ran a linear mixed-effects model on all participants, with Block-Type (tVNS, SHAM) and Trial-History (post-correct, post-error) as within-subject factors. Since pupil size was normalized using z-scores, the absolute z value of pupil baseline is not meaningful, and Group was therefore not included as a factor in this analysis. Then, for pupil dilation, we first used paired t-tests (Block-Type: tVNS, SHAM) with cluster-based Monte-Carlo permutations (1000 iterations), identifying periods of significant differences between tVNS and SHAM blocks for each group. This was done both on the 11-second window aligned to stimulation onset and on the 5-second window aligned to coherent dot motion onset. After that, to compare the overall impact of tVNS on pupil dilation depending on its application time during the task, we computed the area under the curve (AUC) between the tVNS and SHAM pupil response traces (Pupil-AUC Δ_BT_) aligned to stimulation onset, over the whole period of significant differences in the three groups. We also computed the Pupil-AUC Δ_BT_ aligned to coherent dot motion onset. In this case, the analysis window for each participant spanned from the coherent dot motion onset to the participant’s average RT to assess group differences in the tVNS effect during the Decision phase. These two pupil response traces measures were analyzed using separate repeated measures ANOVAs with Group (Early, Mid, Late) as a between-subject factor, Trial-History (post correct, post-error) as a within-subject factor, and Accuracy Δ_TH_ in SHAM blocks included as a covariate, consistent with the approach used in the behavior analysis. To assess whether the two Pupil-AUC Δ_BT_ varied across different performance contexts in the three groups, follow-up repeated-measures ANOVAs were conducted with Trial-Type (LowACC, HighACC) and Group (Early, Mid, Late) as factors.

Finally, in a control analysis, we also checked whether behavior (accuracy, RT) naturally varies as a function of pupil dilation during the decision phase. To do so, we considered data in SHAM blocks and used a generalized (or linear) mixed-effects model, with accuracy (or RT) as dependent variable, Group (Early, Mid, Late) as a fixed factor, and pupil dilation during decision-making as a covariable, which was further divided into four bins for post hoc analyses. All post-hoc comparisons used Tukey corrections.

## Discussion

This study investigated the behavioral impact and underlying computational mechanisms of online tVNS-induced increases in LC-NE activity, applied at three distinct time points across groups performing perceptual decisions in a RDM task. tVNS consistently elicited greater pupil dilation than SHAM stimulation across all groups, indicating effective activation of the LC-NE system during the task. Importantly, early stimulation produced the most sustained activation, with tVNS-induced pupil dilation already robust during the Fixation phase but remaining as pronounced as in the other groups during the Decision phase. Behaviorally, tVNS reliably improved decision accuracy without affecting response times. These improvements were most prominent in trials with initially lower accuracy under SHAM, suggesting that tVNS enhanced performance when initial task engagement was suboptimal, likely by facilitating evidence accumulation, as supported by increased drift rate in the DDM analyses. The effect of tVNS on accuracy was quite consistent across groups but nevertheless stronger when tVNS was applied early in the trial.

Expanding on previous findings (Su, Vanvoorden et al. 2025), the present work shows that tVNS improves decision accuracy without affecting RTs, supporting a gain modulation function for LC-NE (Aston-Jones and Cohen 2005, Devilbiss 2019). Moreover, by applying tVNS at different phases of the decision process in a larger sample (n = 62), this study strengthens evidence for the robustness of the observed effect and demonstrates its generalizability across stimulation timings. Interestingly, our current findings also strongly support the view that tVNS does not uniformly enhance performance across all trials, but instead selectively benefits behavior in trial types that show relatively lower decision accuracy under SHAM. Specifically, for participants who showed reduced accuracy following errors in SHAM blocks, tVNS selectively enhanced performance in post-error trials. Conversely, for those whose accuracy was lower after correct responses under SHAM, performance improvements emerged in post-correct trials selectively. The mechanisms driving these selective effects likely differ between subgroups. Prior studies suggest that when task accuracy is generally high, the occurrence of an error can act as a salient and unexpected signal that captures attention and temporarily disrupts ongoing cognitive control (Notebaert, Houtman et al. 2009, Castellar, Kühn et al. 2010, Houtman, Castellar et al. 2012, Van der Borght, Schevernels et al. 2016). In line with this, participants in our study who showed impaired post-error accuracy had relatively high accuracy in post-correct trials (mean accuracy of 79%), suggesting that errors were infrequent and may have triggered attentional shifts away from task-relevant information. In contrast, participants who exhibited reduced accuracy after correct responses showed generally lower performance even in post-correct trials (mean accuracy of 69%), a pattern that emerged in the absence of error feedback. Similar behavioral dynamics have been reported in sustained attention tasks, where accuracy gradually declines over sequences of correct responses (Helton and Warm 2008, Langner and Eickhoff 2013). These declines have been linked to a gradual disengagement of attention, possibly due to reduced performance monitoring or increased automatization during repetitive task execution (Thomson, Besner et al. 2015). This mechanism likely underlies the worse performance we observed in post-correct trials in a proportion of subjects, where attention may have faded in the absence of salient events. Altogether, these findings suggest that tVNS selectively enhanced accuracy by reinforcing attentional control during periods of weakened endogenous regulation, whether triggered by distraction from unexpected events or by gradual disengagement in repetitive task contexts.

DDM analyses specifically allowed us to assess the computational mechanisms underlying the effect of tVNS on decision performance. Interestingly, this approach revealed a significant increase in drift rate under tVNS, with none of the other DDM factors being significantly influenced by tVNS. Moreover, the effect on drift rate was specific to the low accuracy context, thus paralleling the observation made on accuracy. Specifically, in the SHAM condition, drift rates were lower in low accuracy than high accuracy setting, indicating that evidence accumulation was less efficient in states of poorer performance (Deakin, Schofield et al. 2024). Then, tVNS restored the drift rate selectively in the low accuracy context, bringing it to similar levels as in the high accuracy context, probably causing the accuracy boost in this same condition. The efficiency of evidence accumulation is known to depend on the degree of cognitive engagement. For instance, prior studies have shown that enhanced attentional focus on task-relevant features leads to faster and more reliable accumulation of sensory evidence, as indexed by higher drift rate (White, Ratcliff et al. 2011, Nunez, Vandekerckhove et al. 2017). From this perspective, our findings suggest that tVNS-induced increases in LC-NE activity allowed to stabilize attentional engagement when it was weakened, thereby facilitating more effective accumulation of coherent dot motion direction.

Intriguingly, tVNS induced similar pupil dilation in low and high accuracy contexts, despite its selective impact on behavior in the former condition. This dissociation implies that similar activations of the LC-NE system had distinct behavioral effects. In other words, the selectivity of the tVNS effect on behavior was more likely related to how LC-NE activity interacted with its target areas, than to how tVNS altered LC-NE activity itself. More precisely, tVNS-induced NE release may have only impacted local circuits onto which LC neurons project when the circuit (e.g. visual cortex) operated in a suboptimal way, consistent with the idea that LC-NE allocates cognitive resources depending on internal demands (Mather and Sutherland 2011, Mather, Clewett et al. 2016, Kaduk, Henry et al. 2023). When attentional engagement is already high and neural gain is optimized in task-relevant processes, additional LC-NE activation may offer limited behavioral benefit and could even impose unnecessary metabolic costs (Nassar, Rumsey et al. 2012). In contrast, under conditions of weakened endogenous control, such as attentional lapses or low task engagement, tVNS-induced LC-NE activation may restore effective processing by stabilizing attention and enhancing the integration of task-relevant information. Within this framework, tVNS does not exert a uniform effect on performance but operates in a state-dependent manner, selectively enhancing behavior when internal control alone is insufficient to support optimal decision-making.

A striking aspect of our findings is the strong convergence of findings across the three groups. Although tVNS was delivered at different time points within the task, corresponding to different stages of LC-NE engagement, all groups showed remarkably similar behavioral effects. These were characterized by consistent improvements in decision accuracy without accompanying changes in RTs, and by a shared pattern of selective enhancement in trials associated with lower SHAM performance. The consistency of these effects across stimulation timings suggests that the behavioral efficacy of tVNS does not critically depend on the precise moment of stimulation. Rather, it appears to be shaped by how stimulation interacts with the cognitive dynamics of the task. Prior theoretical and empirical work has shown that the LC does not respond exclusively to external stimuli, but dynamically adjusts its activity based on ongoing task demands and internal cognitive state (Aston-Jones and Cohen 2005, Sara and Bouret 2012, Bouret and Richmond 2015). This context-sensitive modulation enables the LC-NE system to influence behavior over sustained periods of task engagement rather than being limited to transient sensory-driven responses. Supporting this view, neurophysiological evidence has characterized the LC as a differentiated neuromodulatory system capable of sustaining task-dependent modulation over extended timescale (Totah, Neves et al. 2018). Recordings in non-human primates further demonstrate that noradrenergic neurons in the LC encode transitions in cognitive state and predict behavioral re-engagement following errors, highlighting their role in tracking internal task dynamics (Jahn, Varazzani et al. 2020). Within this framework, the convergence of behavioral effects across groups suggests that the impact of tVNS is not determined by when stimulation begins, but by whether it coincides with periods of sustained task engagement, during which LC-NE activity is actively recruited and temporally extended. Under such conditions, tVNS may amplify ongoing LC-NE dynamics that are already tuned to task structure, allowing its effects to persist across multiple phases of the decision process regardless of stimulation onset.

While the behavioral effects of tVNS were broadly consistent across stimulation groups, participants in the Group_Early_ exhibited the most pronounced overall improvement in decision accuracy across the RT spectrum, as reflected by a greater upward shift in the conditional accuracy curve. This enhancement was accompanied by the most sustained pupil dilation following stimulation, with Group_Early_ showing the longest continuous period of pupil enlargement and the highest overall pupil dilation aligned to stimulation onset. These findings suggest that tVNS delivered during the early Fixation phase, when LC-NE activity was still at baseline at the stimulation onset, may be particularly effective, potentially due to its alignment with the onset of task engagement. Prior research indicates that the efficacy of noradrenergic modulation is shaped by the cognitive context in which it occurs, especially during periods when attention is being allocated and control processes are being established (Borodovitsyna, Flamini et al. 2017). Delivering stimulation at the beginning of the task engagement, when internal control systems are actively being configured to support upcoming goal-directed behavior, may offer a privileged window for modulating LC-NE activity. Enhancing LC-NE engagement during this preparatory phase may facilitate the formation of a stable cognitive state and promote sustained attentional allocation throughout the trial. Such temporal alignment between tVNS and early-stage task engagement may allow the resulting neuromodulatory effects to persist across the decision process, thereby contributing to a larger improvement in performance.

## Conclusion

This study demonstrates that tVNS enhances perceptual decision-making by selectively improving decision accuracy without compromising response speed. This enhancement was robust across different stimulation timings and was most pronounced in trials with initially lower accuracy. DDM revealed that tVNS increased the efficiency of evidence accumulation, as reflected by higher drift rates compared to the SHAM condition, without altering decision thresholds or non-decision time. Notably, although tVNS consistently elicited pupil-linked LC-NE activation across trial types, behavioral benefits emerged selectively under conditions of diminished endogenous control. This dissociation suggests that the cognitive impact of tVNS depends critically on the cognitive context in the task, with the LC-NE system exerting its greatest influence when attentional engagement is suboptimal. Moreover, while the overall accuracy enhancement under tVNS was comparable across stimulation timings, early stimulation produced a greater accuracy benefit across the RT spectrum and was accompanied by a stronger pupil response aligned to stimulation onset, indicating enhanced modulation of both behavioral and physiological markers of decision-making. Together, these findings provide compelling causal evidence for the role of the LC-NE system in supporting adaptive decision-making and establish tVNS as a powerful non-invasive tool for modulating cognitive performance in a state-dependent manner. Future work should further explore how task demands interact with neuromodulatory interventions to shape behavioral outcomes.

## Acknowledgments

SS was supported by grants from UCLouvain (FSR mobility grant: SW/AS/17846.2023) and from the Belgian National Fund for Scientific Research (FNRS: FC 48815). The research was also supported by an ARC grant from UCLouvain (Coaction) and research credits from FNRS (PDR UrgeToAct: T007023F and CDR-AROUSAT: J007424F).

